# pH-Sensitive Optical Nanocomposites Using Polymer-Coated Gold Nanorods

**DOI:** 10.1101/2025.03.31.646487

**Authors:** Yongping Chen, Xingde Li

## Abstract

This study investigates the synthesis of pH-responsive, reversible nanocomposites comprising polystyrene sulfonate (PSS)-coated gold nanorods and poly(allylamine hydrochloride) (PAH)-coated gold nanorods, along with their optical properties. We observed a pH-dependent swelling/shrinking of the nanocomposites and a dramatic red-shift (∼ 60 nm) of the surface plasmon resonance (SPR) peaks as the pH changed from around 5.4 to 7.2 due to the increased side-by-side interactions of adjacent gold nanorods. These pH-responsive nanocomposites, with tunable SPR peaks, hold potential for use as contrast agents in optical molecular imaging.

**GRAPHICAL ABSTRACT:** pH-Sensitive Polymer-Coated Gold Nanorods for Reversible SPR Shifts and Applications in pH Sensing as Optical Materials.

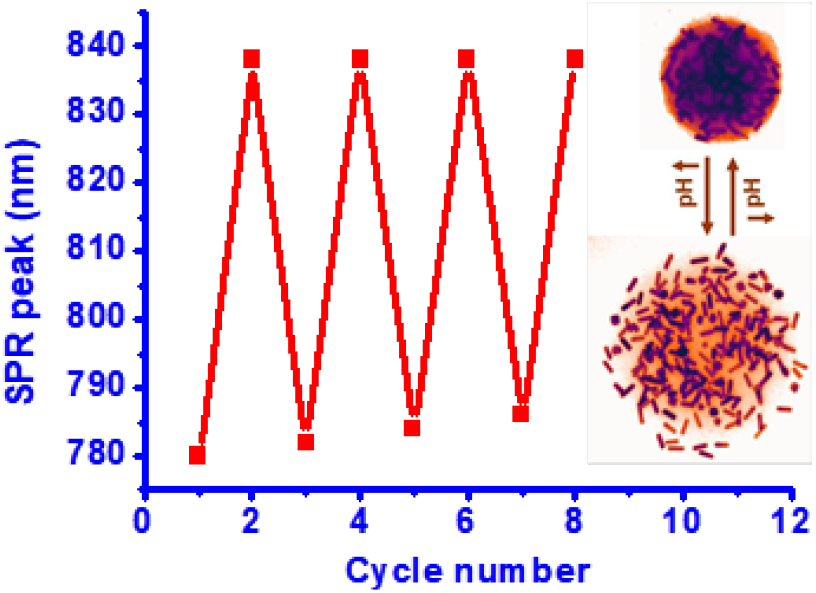

## INTRODUCTION

Gold nanorods (GNRs) have garnered significant interest in recent years due to their distinctive optical properties, which make them promising for various applications, including optical contrast imaging [1-3], drug release [4, 5], photothermal therapy [6, 7], spectroscopy enhancement [8, 9], and optical sensing [10, 11]. These optical properties arise from localized surface plasmon resonance (SPR), which causes characteristic absorption peaks in the visible to near-infrared region, known as plasmon bands [12]. The intensity and wavelength of these surface plasmons are highly sensitive to the dielectric constant of the surrounding medium, and this sensitivity can be leveraged for sensing applications through shifts in SPR peaks [13]. In addition to their optical properties, the presence of positively charged cetyltrimethylammonium bromide (CTAB) molecules (used for nanorod synthesis) on the surface of gold nanorods serves as a stabilizing agent, facilitating straightforward surface modification of the nanorods for further applications [14].

Due to the unique properties of gold nanorods and their potential optical response to external stimuli, there has been growing interest in developing stimuli-responsive systems by incorporating polymers to the surface of nanorods. For instance, Ivan *et al*. developed hybrid microgels composed of a temperature-responsive copolymer and gold nanorods, enabling photothermally modulated volume transitions in the near-infrared (near-IR) spectral range [15]. Similarly, Atsushi Shiotani and colleagues prepared gold nanorod-embedded N-isopropyl acrylamide (NIPAM) hydrogels and investigated their volume phase transition behavior upon near-IR laser irradiation [16]. In both cases, the stimuli-responsive systems, which combined gold nanorods with temperature-sensitive polymers, were designed to explore thermoresponsive optical properties. Matthias and collaborators developed multiresponsive hybrid colloids based on polyelectrolyte-coated gold nanorods and poly (NIPAM-co-allylacetic acid) microgels to study temperature and pH-tunable plasmon resonance at pH 8 and pH10 [17]. However, to date, there have been no reports investigating the development of pH-sensitive SPR peaks in polymer-coated gold nanorod nanocapsules under physiological pH conditions.

In this report, we introduce a class of nanocomposites which were made of gold nanorods coated with poly styrene sulfonate (PSS) and poly allylamine hydrochloride (PAH), and investigated the optical properties of these nanocomposites under physiological pH conditions. The results showed pH-induced swelling/shrinking behavior of the nanocomposite, leading to a pH-responsive and reversible SPR change under physiological pH conditions.

## RESULTS AND DISCUSSION

In this paper, gold nanorods were synthesized using a wet chemical protocol, with CTAB serving as a stabilizing agent [14]. After purification, the gold nanorods remained protected by CTAB, which imparts a strong positive charge to their surface. To manipulate the surface charge, we applied a layer-by-layer coating technique with poly(styrene sulfonate) (PSS), a polyanion, which resulted in nanorods with varying surface charges, from highly positive to highly negative (referred to as type I: PSS@gold nanorods). Subsequently, half of the type I gold nanorods were coated with poly(allylamine hydrochloride) (PAH), a polycation, to yield nanorods with a positive charge on their surface (referred to as type II: PAH/PSS@gold nanorods). Figure 1 illustrates the layer-by-layer assembly process of polyelectrolytes on the nanorod surface (Figure 1a), as well as the structures of PSS and PAH (Figure 1b). Following the PSS coating, the absorption peak of the gold nanorods exhibited a 2-nm red shift in the longitudinal SPR band (from 757 nm). After the PAH coating, a further 4-nm red shift in the SPR was observed relative to the PSS@gold nanorods. Additionally, the absorption spectra of the gold nanorods did not show significant broadening (similar to the one reported by Wang *et al*. [18]), indicating no pronounced particle aggregation (Figure 1c).

**Figure 1.**
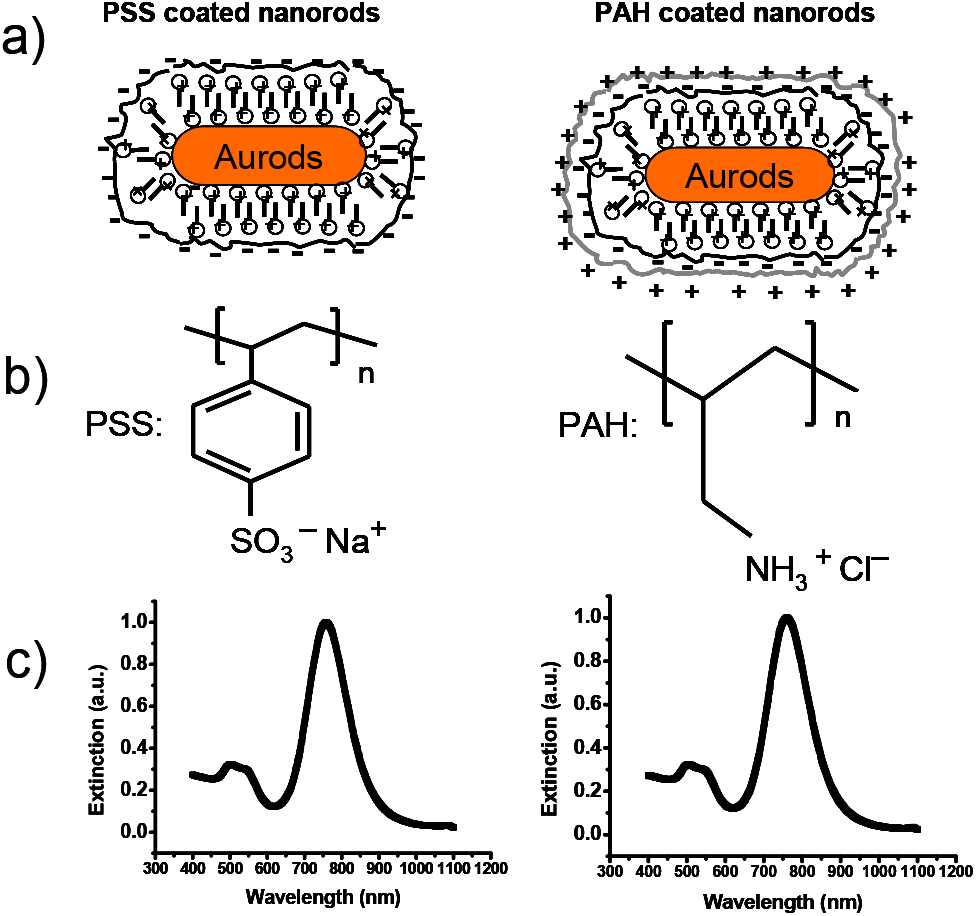
a) Schematic representation of the layer-by-layer process for preparing two types of gold nanorods: PSS-coated gold nanorods and PAH/PSS-coated gold nanorods. b) Structures of PSS and PAH. c) Extinction spectra of PSS-coated gold nanorods (peak at 757 nm) and PAH/PSS-coated gold nanorods (peak at 761 nm).

For the formation of nanocomposites, 0.5 mL of PSS-coated gold nanorods (1 nM) was mixed with an equal volume of PAH-coated gold nanorods in an aqueous solution. Following the mixing, electrostatic interactions between the PSS-GNs and PAH-GNs induced their aggregation, leading to an 8-nm red shift in the SPR peak (from 761 nm to 769 nm) (Figure 2a). However, this interaction did not result in nanoparticle aggregation, as observed in the TEM images (Figure 2b). Given that PAH and PSS are sensitive to pH, we added NaOH to trigger stronger interactions between the PSS-GNs and PAH-GNs. Upon the addition of NaOH, a rapid red shift occurred, and the absorption spectra of the gold nanorods exhibited notable broadening, indicative of nanoparticle aggregation. After approximately 60 minutes, the surface plasmon resonance peak shift reached its maximum. For a 1 mL mixture of gold nanorods with the addition of 7 μL of 0.1 M NaOH (resulting in a solution pH of ∼9.3), the red shift extended from 769 nm to 838 nm (Figure 2c), and TEM images showed the aggregation of gold nanorods due to self-assembly between the PSS-GNs and PAH-GNs (Figure 2d).

**Figure 2.**
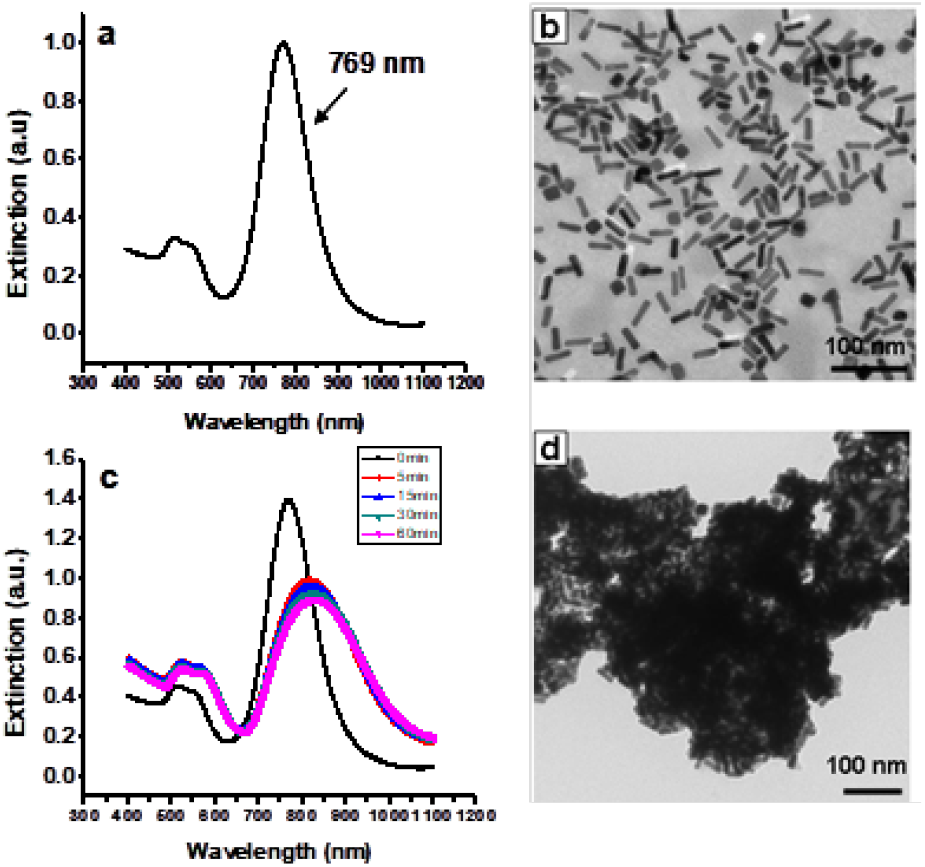
a) Absorption spectra of the mixture of PSS-GNs and PAH-GNs. b) TEM images of the mixture of PSS-GNs and PAH-GNs, showing no aggregation. c) Dynamic changes in the extinction spectrum. d) TEM images of the mixture solution of PSS-GNs and PAH-GNs, showing aggregation after the addition of NaOH.

Although PSS-GNs and PAH-GNs can self-assemble in basic solutions, the resulting polyelectrolyte complexes do not form a regular structure (Figure 2d). The presence of NaCl in polyelectrolyte solutions significantly affects the chain configuration in the bulk solution. In the absence of salt, the same charges within a single polyelectrolyte chain repel each other, causing the chain to adopt an almost fully extended, rod-like configuration. However, in the presence of salt, counterions screen some of the charges, allowing the polyelectrolyte chain to fold into a random coil configuration [19]. We induced the self-assembly of PSS-GNs and PAH-GNs in a salt solution under two different pH conditions (pH 9.30 and pH 9.8). At both pH conditions, spherical assemblies were observed. TEM images revealed the formation of nanocomposites with a diameter of 254 ± 86 nm at pH 9.30 (referred to as type I nanocomposite), where each gold nanorod was clearly visible, as shown in the inset (Figure 3a). At pH 9.8, the nanocomposites had a larger diameter of 306 ± 106 nm (referred to as type II nanocomposite) (Figure 3c). The larger assemblies appeared more solid, and the individual gold nanorods constituting the sphere were less distinguishable. Due to their larger size and packing, the number of nanorods in the type II nanocomposite was higher than in the type I nanocomposite. This higher number of gold nanorods in the type II nanocomposite led to a larger red shift in the SPR peak. The UV-vis spectrum showed a red shift from 769 nm to 834 nm for the type I nanocomposite (Figure 4a) and a red shift from 769 nm to 864 nm for the type II nanocomposite (Figure 4c). Since the degree of ionization of the polymer is pH-sensitive, the properties of the nanocomposites can be tuned by changing the pH. When 0.1 M HCl was added to 1 mL of the type I and type II solutions, the SPR wavelength of the type I nanocomposites shifted from 834 nm to 780 nm, and the type II nanocomposites shifted from 864 nm to 789 nm, with the final solution pH being approximately 5.4. TEM images showed the swelling of the nanocomposites, with a diameter of 457 ± 124 nm (pH ∼ 5.4) for type I nanocomposites (Figure 3b), and 654 ± 316 nm (pH ∼ 5.4) for type II nanocomposites (Figure 3d).

**Figure 3.**
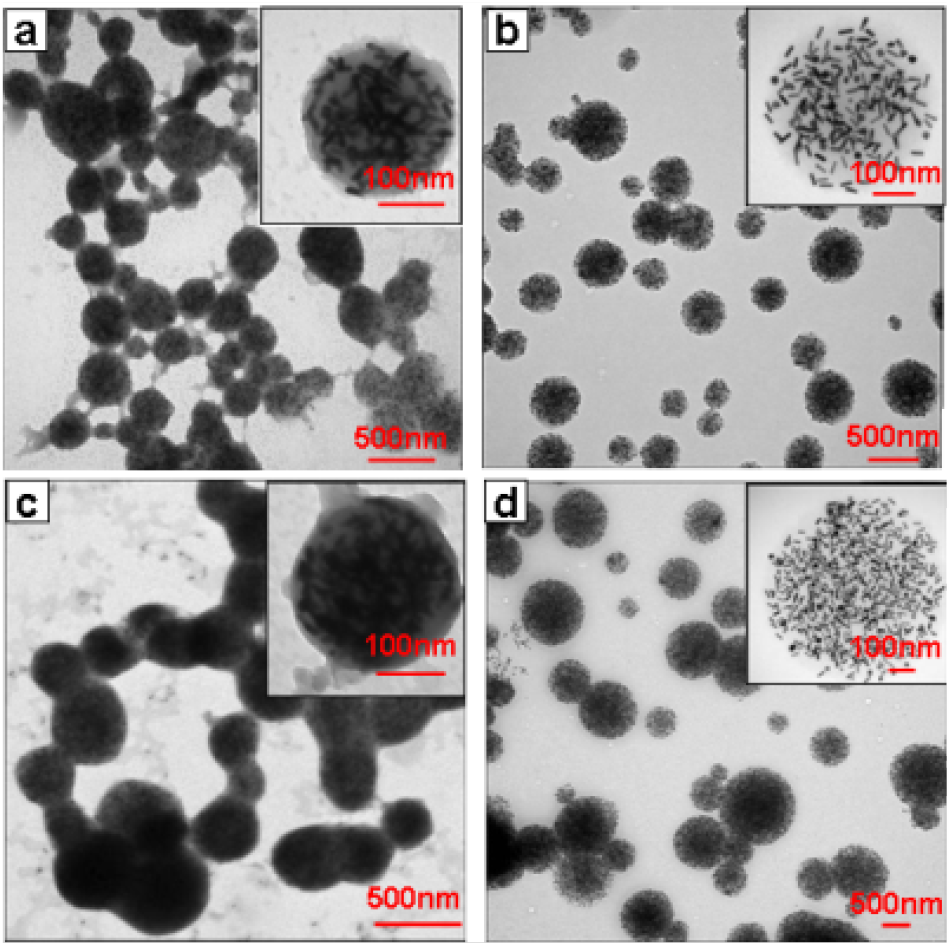
TEM images of Type I nanocomposites with a diameter of 254 ± 86 nm at pH 9.30 (a) and their swelling at pH 5.4, resulting in a diameter of 457 ± 124 nm (b). TEM images of Type II nanocomposites with a diameter of 306 ± 106 nm at pH 9.8 (c) and their swelling at pH 5.4, resulting in a diameter of 654 ± 316 nm (d).

**Figure 4.**
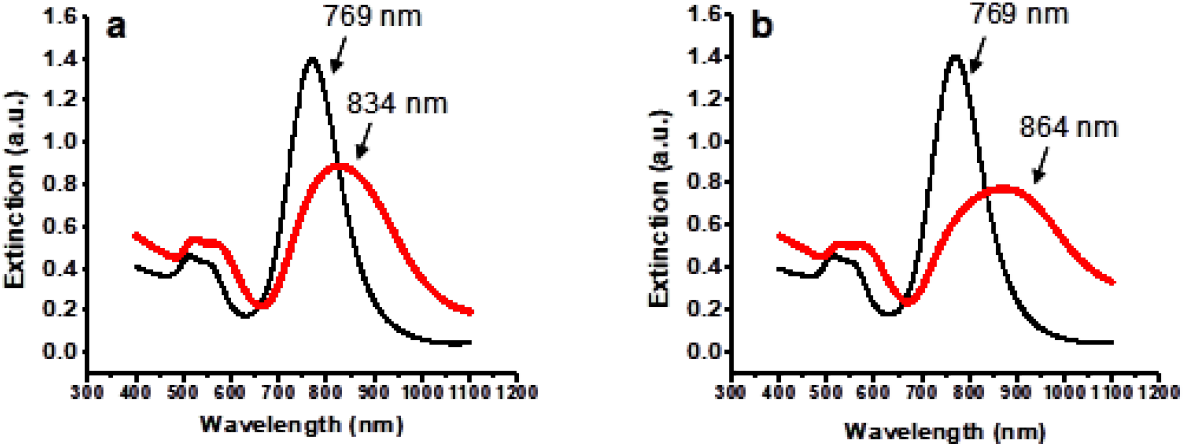
The UV-vis spectrum shows an SPR red-shift from 769 nm to 834 nm at pH 9.30 (a) and an SPR red-shift from 769 nm to 864 nm at pH 9.80 (c).

The behavior of shrinking and swelling in the nanocomposites with pH changes (Figure 3) demonstrates the potential to tune the SPR peak of gold nanorods by varying the pH. To investigate the pH-responsive properties of the nanocomposites synthesized in this study, the SPR peaks of the nanocomposites were measured under various physiological pH conditions (ranging from 5.0 to 7.4). Figure 5a shows the relationship between pH and the SPR peak of the two types of nanocomposites, as determined from UV-vis spectra under physiological pH conditions. It was observed that the SPR peak of type I nanocomposites shifted from 780 nm at pH 5.4 to 838 nm at pH 7.2, while the SPR peak of type II nanocomposites shifted from 789 nm at pH 5.4 to 868 nm at pH 7.2. Additionally, the pH-dependent shifts in the SPR peaks were reversible, as shown in Figure 5b. Alternating the pH between 5.4 and 7.2 resulted in a reversible change in the SPR peak from approximately 780 to 838 nm for type I nanocomposites and from approximately 789 to 868 nm for type II nanocomposites.

**Figure 5.**
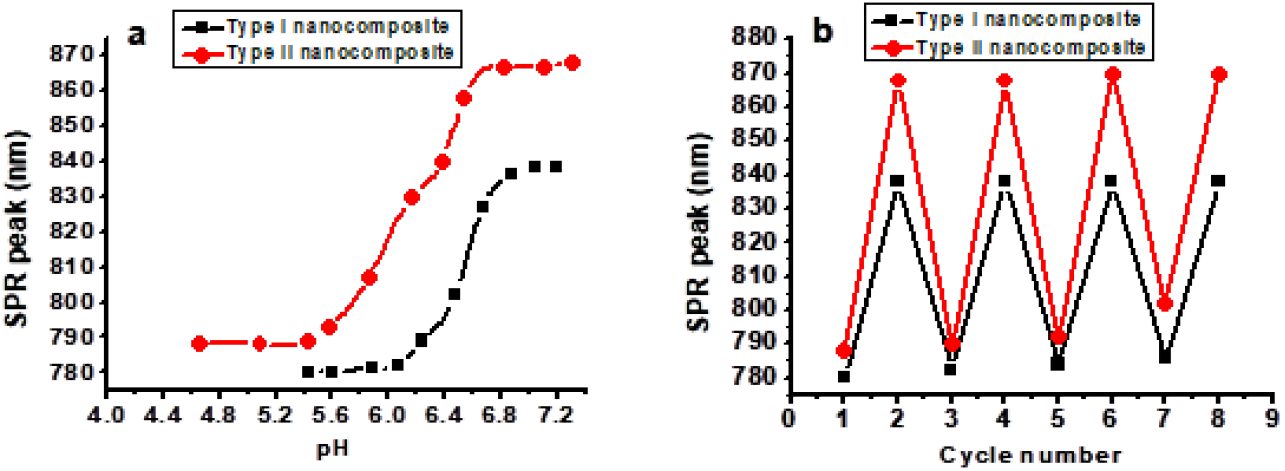
The SPR peaks of nanocomposites are sensitive to pH changes under physiologic pH conditions (a), and the changes are reversible (b).

The data obtained can be summarized and explained as shown in Figure 6. At neutral pH, the polymer on the surface of the gold nanorods exhibits maximum stiffness, and electrostatic interactions are insufficient to overcome the orientation entropy of the polymer, preventing aggregation of the gold nanoparticles (Figure 6a) [20]. NaCl salt plays a key role as a physico-chemical parameter, regulating the strength of ionic bonding, the conformation of the polymer (from linear to coil configuration), and its mechanical properties (from stiffness to softness) [19]. Therefore, in NaCl solution, NaOH can trigger the self-assembly of PSS-GNs and PAH-GNs into nanocomposites through electrostatic interactions between the polymer-coated nanoparticles and hydrophobic interactions in PSS (Figure 6b) [21]. After synthesizing the nanocomposites in basic NaCl solution, the polymers undergo irreversible interactions, and a decrease in pH does not induce polymer desorption. However, charging of the PAH layer may occur, introducing a positive charge into the nanocomposites. This change may alter the morphology of the nanocomposites by enhancing mutual repulsion, which could lead to the swelling of the nanocomposites under acidic conditions (Figure 6c) [22]. Consequently, the SPR peaks of the nanocomposites can be tuned, causing swelling (Figure 6c) and shrinking (Figure 6d) by varying the pH within the physiological range.

**Figure 6.**
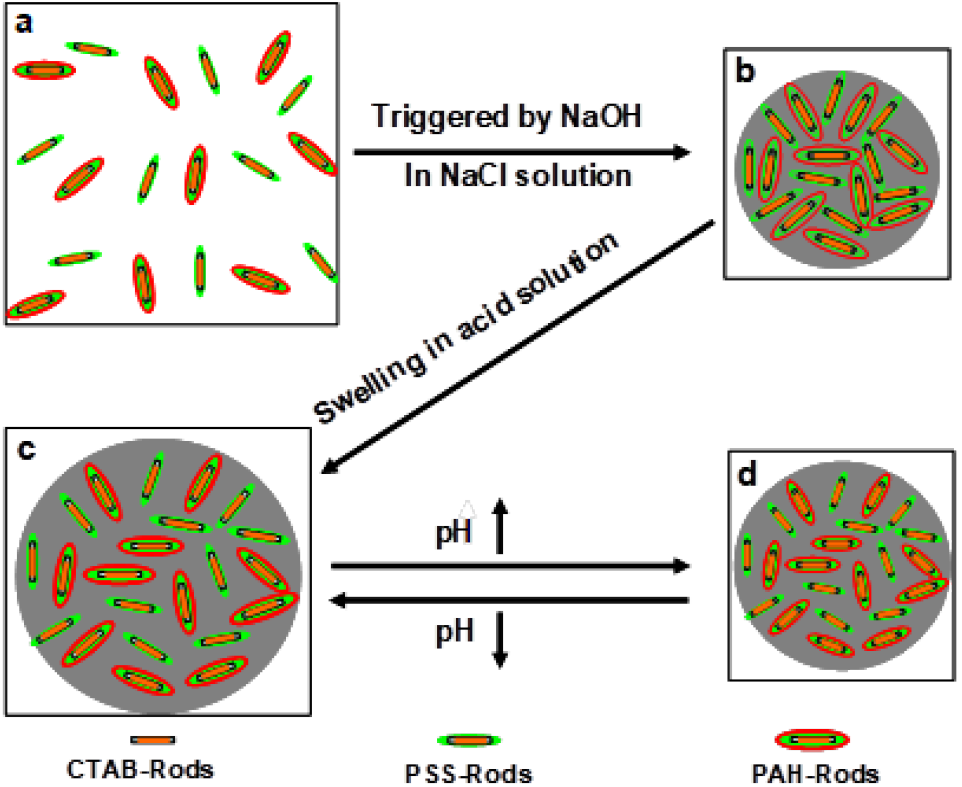
Model of the synthesis of nanocomposites and tuning of SPR peaks in nanocomposites by pH changes.

## CONCULIONS

We report on the development of pH-sensitive nanocomposites with a broad surface plasmon resonance (SPR) tuning range. These nanocomposites are composed of polymer-coated gold nanorods (PSS @gold nanorods and PAH/PSS @gold nanorods) in NaCl solution, with pH changes triggered by NaOH. The size of the nanocomposites can be finely tuned by adjusting the solution pH. The resulting SPR peak shift is highly sensitive and reversible, allowing for precise tuning within the physiological pH range, which leads to notable shifts in the SPR peaks. These nanocomposites are promising for pH-sensing applications and hold potential as contrast agents for optical imaging. To the best of our knowledge, this is the first report on the fabrication of pH-sensitive nanocomposites through the polyelectrolyte coating of gold nanorods, as well as an investigation into the optical properties of these nanocomposites.

## EXPERIMENT SECTION

### Synthesis of Gold Nanorods

Gold nanorods were synthesized using the seed-mediated method [14, 23]. To prepare the gold seeds, 0.1 mL of a 0.01 M HauCl_4_·3_2_ O aqueous solution was injected into 3 mL of a 0.1 M CTAB aqueous solution and mixed by inversion. Then, 0.24 mL of ice-cold 0.01 M NaBH4 was added, followed by rapid inversion mixing for 2 minutes. This seed solution was used 2 hours after its preparation and remained usable for up to two days.

### Preparation of Gold Nanorods

A solution containing 42 mL of 0.1 M CTAB was mixed with 0.45 mL of 0.01 M silver nitrate aqueous solution, 1.68 mL of 0.01 M HauCl_4_·3H_2_O aqueous solution, and 0.28 mL of 0.10 M L-ascorbic acid aqueous solution, with continuous stirring. Subsequently, 0.28 mL of the seed solution was injected into the growth solution. After shaking for 10 seconds, the mixture was left undisturbed for 24 hours at room temperature.

### Purification of the Gold Nanorods

After 24 hours of growth, the solution was filtered using filter paper and then centrifuged at 8000 rpm at room temperature to remove excess CTAB and other reagents. Finally, the deposited gold nanorods were redispersed in 20 mL of pure water for storage.

### Polyelectrolyte-Coated Gold Nanorods

The layer-by-layer modification of gold nanorods has been reported previously [24]. In this work, a 3 mg/mL concentration of the polyelectrolytes PSS and PAH with 6 mM NaCl was used. First, 20 mL of the gold nanorods solution was added drop by drop to an equal volume of PSS under sonication. After adding the gold nanorods, the mixture was stirred overnight. Next, the solution was centrifuged at 8000 rpm, the supernatant was discarded, and the gold nanorods were re-dispersed in 20 mL of water. Subsequently, 10 mL of the PSS-coated gold nanorods was added dropwise to 10 mL of the PAH solution under sonication, stirred for 12 hours, centrifuged, and stored in 10 mL of Milli-Q water. The coated nanorods remained stable for weeks, as indicated by the absence of any color changes over time.

### Synthesis of Nanocomposites by the Polymer-Coated Gold Nanorods

0.25 mL of PSS-coated gold nanorods were diluted in 0.5 mL of Milli-Q water with 12 mM NaCl (final concentration of gold nanorods ∼ 1 nM). Next, 0.25 mL of PAH-coated gold nanorods was diluted to 0.5 mL with pure water (final concentration of gold nanorods ∼ 1 nM) and added dropwise to the above solution under stirring. Then, either 7 μL or 8.5 μL of 0.1 M NaOH aqueous solution was added, followed by stirring for 20 seconds. The color was observed to change over time, from red wine to purple. The reaction stabilized in about 1 hour. During the experiments, UV-vis spectroscopy was performed using a DU series 700 Scan UV-vis spectrophotometer from Beckman Coulter Inc., pH measurements were taken using a portable Chekmite pH-20, and transmission electron microscopy (TEM) measurements were conducted with a Hitachi H-7000 TEM instrument operating at an accelerating voltage of 200 kV. For TEM, samples were prepared by placing a 3 μL drop of the purified gold nanorod solution on Formvar-coated copper grids (200 meshes, Ted Pella Inc.) and drying for 1 hour.

## Acknowledgements of Funding Supports

This work was supported in part by the National Institutes of Health (NIH) (R01 CA120480 and 1R01 EB007636)

